# Population heterogeneity in mutation rate increases mean fitness and the frequency of higher order mutants

**DOI:** 10.1101/045377

**Authors:** Helen K. Alexander, Stephanie I. Mayer, Sebastian Bonhoeffer

## Abstract

Mutation rate is a crucial evolutionary parameter that has typically been treated as a constant in population genetic analyses. However, mutation rate is likely to vary among co-existing individuals within a population, due to genetic polymorphisms, heterogeneous environmental influences, and random physiological fluctuations. We explore the consequences of such mutation rate heterogeneity in a model allowing an arbitrary distribution of mutation rate among individuals, either with or without inheritance. We find that variation of mutation rate about the mean results in a higher probability of producing zero or many simultaneous mutations on a genome. Moreover, it increases the frequency of higher order mutants even under ongoing mutation and selection. We gain a quantitative understanding of how this frequency depends on moments of the mutation rate distribution and selection coefficients. In particular, in a two-locus model, heterogeneity leads to a relative increase in double mutant frequency proportional to the squared coefficient of variation of the mutation rate. Relative effect sizes increase with the number of loci. Finally, this clustering of deleterious mutations into fewer individuals results in a higher population mean fitness. Our results imply that mutation rate heterogeneity allows a population to maintain a higher level of adaptedness to its current environment, while simultaneously harboring greater genetic diversity in the standing variation, which could be crucial for future adaptation to a new environment. Our results also have implications for interpreting mutation rate estimates and mutant frequencies in data.

## 1 Introduction

The mutation rate is a key evolutionary parameter that affects the level of genetic diversity in a population. Genetic diversity in turn affects both the population’s current mean fitness and its capacity to adapt to changes in the environment. Most theoretical work to date has assumed that the mutation rate takes on a fixed value in all members of the population. Nonetheless, the mutation rate, like any other trait, can be expected to vary among individuals, due to genetic, environmental, and stochastic effects. The recognition that mutation rate can vary within a population is implicit in the long-standing study of mutation rate evolution, and more recently in considerations of transient or stress-induced mutagenesis, especially in bacteria. However, a comprehensive conceptual understanding of how mutation rate heterogeneity within a population affects the level of standing genetic variation is lacking.

The existence of rare individuals with high mutation rate could be particularly important when a combination of several mutations is relevant for adaptation [62, 8, 23, 22]. Given that the mutation rate is typically low, higher order mutants are generally rare, yet they can be crucial for adaptation to complex new environments. For example, when multiple drugs are applied in combination – a common treatment approach for cancer [1] and several major infectious diseases, including tuberculosis, malaria, and HIV [32] – resistance in the targeted pathogens/cells generally requires multiple mutations. The prevalence of such mutations in the “pre-existing” or standing genetic variation before drug treatment starts, when they are generally expected to be deleterious, is predicted to be crucial to the emergence of resistance during treatment [67, 41]. Multiple mutations are also involved in the initiation and progression of many cancers [39].

There is clearly a genetic contribution to mutation rate via genes involved in replication, proofreading, and repair of the genetic material. This can result in variation of mutation rate even among closely related individuals. Laboratory investigations have identified “mutators” and “antimutators” (having higher, resp. lower mutation rate than the wild type), attributable to one or few specific genetic changes, in a variety of organisms. Effect sizes range up to hundreds‐ to thousands-fold variation in eukaryotic cells, bacteria and DNA viruses, and up to around five-fold in retro‐ and RNA viruses (details in Supplementary Text I.1).

Though these studies indicate the scope for variation, the abundance of such variants in natural populations is less clear. Mutators are expected to arise frequently *de novo* due to the large target size for mutations causing defects in replication or repair genes [21, 19]. Theoretically, under constant conditions, alleles that alter mutation rate can be expected at mutation-selection balance in the long term [35, 19, 50, 20]. Moreover, by hitchhiking with beneficial alleles they generate during phases of adaptation, mutators may rise to higher frequency in the short term [75, 45, 20]. In experimental populations of bacteria, mutators have indeed been observed to spontaneously arise and persist [73] and be enriched through selective sweeps [55], and some parameters of these processes can be estimated [9]. Surveys of clinical and other natural isolates in several bacterial species indicate that strains exhibiting a range of mutation rates also exist outside the laboratory [42, 56, 63, 7, 18, 68, 59, 64, 3, 78]. In RNA virus populations, mutators appear rapidly in laboratory settings [74, 13] and are expected to be present in heterogeneous natural populations [74, 52], but we are not aware of any surveys of natural isolates. Cancerous tumors, which are characteristically genetically unstable and highly heterogeneous [44, 31, 4], are also anticipated to be polymorphic in genes affecting mutation rate. It has been hypothesized that a mutator phenotype arises early in carcinogenesis, and moreover increases the chances of successive mutations affecting genomic stability, leading to further non-uniform increases in mutation rate [49, 46, 47, 48]. However, in no case does there appear to be a study quantifying mutation rate in a representative sample of co-existing individuals from a single population (within one infected patient or tumor).

Many environmental factors – including temperature, pH, oxygenation, UV radiation, and chemicals – have also been implicated in modulating mutagenesis in bacteria, viruses, and cancerous cells (details in Suppl. Text I.2). Viral mutation rate could also be affected by its host cell’s type, physiological state, and antiviral defenses. However, few quantitative estimates relating environmental variables to mutation rate are available. Some antibiotics appear to increase bacterial mutation rates by 2‐ to around 100-fold [30, 40], while certain antiretrovirals increase HIV-1 mutation rate by roughly five-fold [53]. While it is clear that the relevant environmental factors may be heterogeneously distributed in a population’s habitat, inducing different mutation rates in coexisting individuals, the precise distribution will be highly context-dependent.

Finally, mutation rate may vary randomly and non-systematically in a population, due to stochastic effects on individuals’ physiological states [9, 22]. For example, the SOS response, which is associated with production of error-prone polymerases in bacteria [76], exhibited a distribution of induction levels in wild type *E. coli* K12, including 0.3% of the population at least 20-fold above the average level at a given time [57]. Even constitutively expressed replication/repair genes are subject to random errors in transcription and translation that affect the protein’s fidelity [62, 8, 58]. Rough calculations suggested that bacterial populations contain resulting “transient mutators” at a total frequency of around 5 × 10^‒4^, with mutation rates expected to be enhanced to similar degrees as in genetic mutators [62]. Fluctuations in low copy number proteins, particularly upon cell division, could also yield temporary reduction in repair capacity [22], and imbalanced concentrations of protein subunits could produce polymerases missing the proofreading subunit [2]. Thus, even isogenic populations in uniform macroenvironments seem likely to contain individuals with differing propensities to generate mutations, although the few tests to date have yielded mixed results [27, 37].

Taken together, this evidence suggests that mutation rate variation within populations is probably common, though there are few direct quantitative estimates. DNA-based organisms appear to have the capacity to vary mutation rate over a few orders of magnitude, while RNA-based viruses appear to tolerate only modest (up to around five-fold) changes in their already high baseline mutation rates [25, 52]. The frequency of mutators in a population could vary widely depending on the source of mutation rate variation and the selective conditions. Furthermore, a broad spectrum arises in the extent to which mutation rate is potentially correlated between parent and offspring. At one extreme, if mutation rate is entirely genetically controlled, the offspring will inherit its parent’s mutation rate. At the other extreme, erroneously translated polymerases or other intracellular components will have limited if any inter-generational effects before degrading and/or being diluted by new production. If mutation rate is primarily determined by the external environment, parent-offspring correlation could vary over a broad range, depending on the extent to which they share a common environment. If spatial variation in the environment is fine-grained (relative to the typical offspring dispersal distance), correlation will be low, while if variation is coarse-grained, parent and offspring are likely to experience the same environment and thus mutation rate.

A large body of theoretical work on evolution of mutation rate takes into account the existence of heritable mutation rate variants (reviewed by [72]). Though the majority of this literature assumes the mutation rate is constitutive, evolution of stress-induced mutagenesis has also been considered [6, 65, 66]. A key factor considered to drive evolution of mutation rate is indirect selection through linkage to other loci that affect fitness, but the focus of these studies is on the dynamics of the mutator allele itself. Far fewer studies have considered how the existence of mutation rate variability in the population, regardless of its source, affects mutational dynamics at other loci [62, 29, 11, 2, 33, 50]. Moreover, these existing models are mostly designed for particular populations and mechanisms of variation, and allow only two possible values of mutation rate.

In the present study, we develop a more general theoretical framework to understand the effects of population heterogeneity in mutation rate on the appearance of new genetic variants at one or more loci, and, in conjunction with fitness, the long-term frequency at which these variants are present. That is, we address not only the production of mutants in a single round of replication, but also the temporal dynamics of deleterious mutants under ongoing production and selection. We consider haploid, asexually reproducing individuals, which is a reasonable first approach for many disease-causing microbial and cellular populations of interest, including bacteria (neglecting horizontal gene transfer in some species), viruses (neglecting complementation and in some cases recombination), and cancerous cells (neglecting dominance effects). Our approach allows an arbitrary distribution of mutation rate among individuals, and considers how moments of this distribution and the degree to which mutation rate is inherited affect the population-level frequency of mutants at focal fitness-determining loci. We do not make any assumption as to the biological mechanism underlying this heterogeneity, in particular whether it is an adaptive/regulated response or an unavoidable byproduct of random processes or external environmental factors.

We find that variability of the mutation rate about the mean has no effect on single point mutants, but boosts the frequency of higher order mutants, with increasingly large relative effects. Through analytical approximations we gain a quantitative understanding, elucidating in particular for a two-locus model that the increase in double mutant frequency is proportional to the variance in mutation rate and depends on the fitness of all mutants. Inheritance of mutation rate strengthens the effect of population heterogeneity, especially when stepwise accumulation of multiple mutations is an important pathway. Finally, we show that a population maintaining a range of mutation rates (whether or not these are inherited) achieves a higher mean fitness than a population in which all individuals have an identical mutation rate.

## 2 Methods

We model a haploid, asexually reproducing population with non-overlapping generations, and extend classic population genetic models to incorporate a mutation rate that varies among co-existing population members. We focus on genotype dynamics at one or more fitness-determining loci (each of which may consist of one or several base pairs), and assume throughout that mutation rate neither depends on the genotype at the focal loci, nor has any direct fitness effect itself.

Data availability R code used to generate numerical results is available upon request.

### 2.1 Occurrence of mutations on a genome in one generation

We consider *n* loci of interest on the genome, which can be either non-mutant or mutant. We assume that each individual has a given mutation rate *u* (per locus, per generation) that is uniform across loci; that is, each non-mutant locus in the individual mutates independently with probability *u*. We neglect back mutations. The number of new mutations that arise thus follows a binomial distribution; in particular, if *n* loci are non-mutant, then the probability of *j* mutations occurring simultaneously (i.e. in one generation) is:

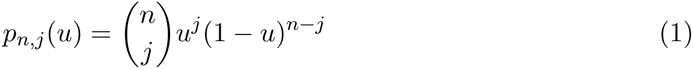

In the limit as *n* → ∞ and *u* → 0 such that *nu* ≡ *λ*, we obtain an “infinite-locus” model in which every mutation occurs at a unique site. Then the number of new mutations that arise in an individual with mutation rate *λ* (per genome, per generation) follows a Poisson distribution; that is, the probability of *j* simultaneous mutations is:

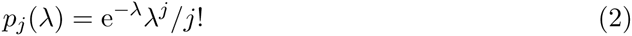

We note that viruses can have complex, multi-step intracellular replication cycles, which imply that one cycle of cell infection cannot be equated to one genome replication, and therefore a Poisson-distributed number of mutations per genome is not necessarily expected after a single infection cycle [26, 71]. Our present model does not address these complexities.

The key novelty in our model is to consider a mutation rate (*u* or *λ*) that varies among individuals in any given generation, and can thus be taken as a random variable (denoted *U* or Λ respectively) in the population as a whole. However, we make the important assumption that the *distribution* of mutation rate in the population does not change over generations.

### 2.2 Inherited versus non-inherited mutation rate

Once we consider dynamics over more than one generation, we must define the extent to which mutation rate is inherited or correlated from parent to offspring. As described in the Introduction, this correlation could vary over a broad spectrum. Mathematically, we will deal with the two extremes.

In the case of no inheritance, each individual in each generation independently draws its mutation rate from the population distribution. Thus one must average the probability or proportion of individuals in the new generation mutating from genotype *i* to *j*, conditioned on mutation rate, over the distribution of mutation rate, which is arbitrary but fixed over generations.

In the case of perfect inheritance, each individual takes on exactly the same mutation rate as its parent. Thus, the population can be divided into subpopulations characterized by distinct mutation rates, with no “migration” among subpopulations. Assuming that the subpopulations do not interact, we can describe the population dynamics in each subpopulation separately using a standard model with fixed mutation rate, before finally taking a weighted average of the quantity of interest over subpopulations. If population size regulation acts on the population as a whole, the subpopulations are not truly independent in their population dynamics. In this situation, indirect selection on mutation rate (due to linkage with focal loci) arises, and if mutations are always deleterious, the lowest mutation rate will be favored [72]. Nonetheless, in line with our aforementioned assumption that the mutation rate distribution does not change over time, we will neglect this predicted evolution of mutation rate, in that we impose selection within each subpopulation independently. Since we will consider selection coefficients at the focal loci that are much larger than any subpopulation’s mutation rate, genotype frequency dynamics at the focal loci are expected to occur on a faster timescale than the evolution of mutation rate (cf. [72, 20]), so our approach should provide a reasonable approximation on this faster timescale.

### 2.3 Genotype frequency dynamics under mutation and selection

We now derive deterministic recursions describing the change in frequency of each genotype from one generation to the next, incorporating both mutation and selection. As a basis, our model adopts a standard formulation in discrete population genetic analyses. Genotypes are defined by the presence/absence of mutations at the focal (fitness-determining) loci. We denote the frequency of genotype *i* at generation *t* by *x*_*i*_(*t*) and its relative fitness by *w*_*i*_. Without loss of generality we take the wild type (carrying no mutations) to have relative fitness 1. We assume that mutations are deleterious, thus *w*_*i*_ = 1 – *s*_*i*_ where the selection coefficients satisfy 0 < *s*_*i*_ *≤* 1 for all types *i* other than the wild type. (Some results below are also valid for the neutral case, *s*_*i*_ = 0, but certain approximations will break down when *s*_*i*_ is too small.) The population mean fitness at time *t* is given by *w̄*(*t*):= ∑_*i*_ *w*_*i*_*x*_*i*_(*t*). Census occurs immediately after mutation, before selection (i.e. relative fitness determines the total reproductive output of parents, offspring mutate independently of one another, then they are counted). Generally, then, for any collection of types *i* and proportions *p*_*ij*_ of type *i* offspring from a parent of type *j*, one can write a set of recursions:

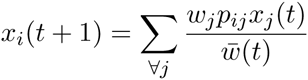

Note that these equations describe the change in genotype frequencies even in a population with changing total size, or absolute fitnesses that change over time, as long as the *relative* fitnesses of the types are constant [16, p. 278]. The incorporation of mutation rate heterogeneity, according to the considerations of sections 2.1 and 2.2, essentially lies in the structuring of the population and in the specification of mutation probabilities {*p*_*ij*_}.

#### 2.3.1 Finite loci

Considering *n* biallelic loci yields *2*^*n*^ types, identified by binary notation indicating absence (0) or presence (1) of a mutation at each locus. For finite *n*, we closely follow the exposition and basic results for fixed mutation rate given by Bürger [10, Ch. III.1.1]. The above recursions can be rewritten as a matrix equation:

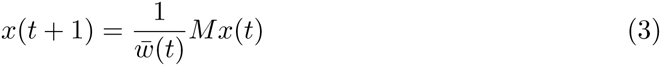

where *x*(*t*) collects the frequencies of each genotype at time *t* into a vector, and *M* is the *2*^*n*^ × *2*^*n*^ “mutation-selection matrix” (independent of time) where *M*_*ij*_ = *W*_*j*_*p*_*ji*_.

Given a mutation-selection matrix *M* and an initial frequency vector *x*(0), the population mean fitness can be written as

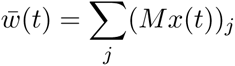

and the solution of the recursion is then given by

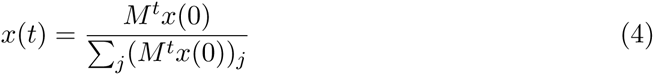

The equilibrium frequency solutions (denoted *x*^*^) are given by the eigenvectors of *M*, normalized so the entries add up to one, and the population mean fitness at equilibrium (*w̄*^*^) is given by the corresponding eigenvalues. Since we neglect back mutation, *M* will always be triangular, and thus the eigenvalues are simply the diagonal entries. The reduction in fitness compared to a homogeneous wild type population due to the production of deleterious mutants, i.e. 1 – *w̄*^*^, is known as the “mutational load” [10, p. 105].

Given the binomial mutation model (Section 2.1), the mutation probabilities between types take the form:

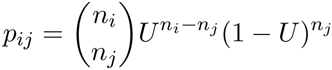

where *U* is the per-locus mutation rate and *n*_*i*_ (resp. *n*_*j*_) is the number of non-mutant loci in type *i* (resp. *j*). We will write the mutation-selection matrix as *M*(*U*) to emphasize its dependence on *U*. For instance for one locus,

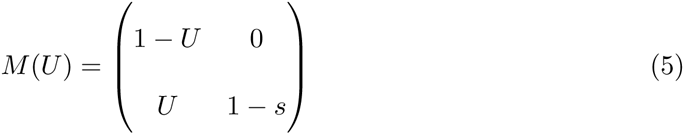

while for two loci,

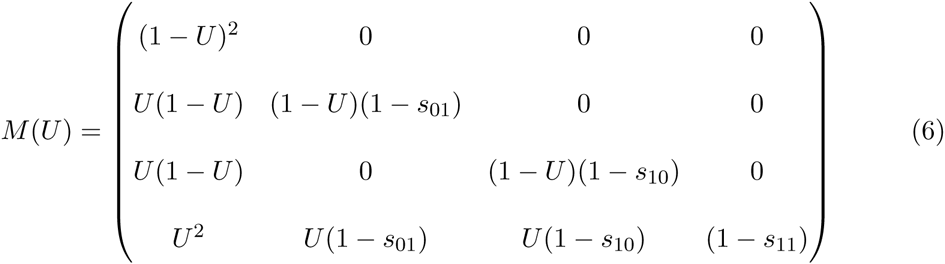

We now incorporate mutation rate heterogeneity at the population level. Although we phrase the following exposition in probabilistic terms, in the present deterministic model we ultimately treat the probability of a type *j* parent producing a type *i* offspring as the exact proportion of such events. As well as neglecting demographic stochasticity, which is standard in this modeling approach, we neglect sampling effects from the underlying mutation rate distribution, which is reasonable if the population is large.

##### Non-inherited mutation rate

Each individual (i.e.each offspring) independently draws its per-locus mutation rate from the distribution of *U*, and conditioned on the mutation rate, the number of non-mutant loci that mutate is binomially distributed (Section 2.1). Applying the Law of Total Expectation, the expected overall contribution of type *j* parents to type *i* offspring is thus given by E_*U*_[*M*_*ij*_(*U*)]*x*_*j*_/*w̄*. This yields the recursion:

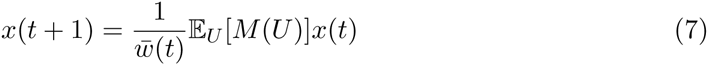

where the expectation over *U* is applied entry-wise to the matrix *M*(*U*).

##### Perfectly inherited mutation rate

The population is divided into *d* disconnected subpopulations, where the *k*^th^ subpopulation is at frequency *q*_*k*_ (with 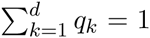) and is characterized by mutation rate *u*_*k*_ that is fixed for all individuals within the subpopulation. Equivalently, the mutation rate distribution in the entire population is given by the probability mass *q*_*k*_ at value *u*_*k*_. Neglecting long-term mutation rate evolution as explained in Section 2.2, we independently solve the recursion in each subpopulation:

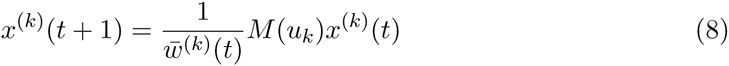

where *x*^(*k*)^ is the genotype frequency vector in the *k*^th^ subpopulation and *w̄*^(*k*)^ (*t*) is the mean relative fitness calculated only within the subpopulation. The population-wide frequencies are finally obtained by averaging over subpopulations, i.e. taking the expectation of the fixed-rate results over the distribution of *U*:

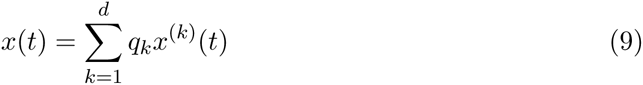

#### 2.3.2 Infinite loci

In the infinite-locus limit, we adopt the model of [38], which assumes that fitness is fully determined by the number of mutations carried. Let *x*_*i*_(*t*) be the frequency of *i*-point mutants in generation *t*; *w*_*i*_ be the relative fitness of *i*-point mutants; and *λ* be the mean number of mutations per genome per generation. The only assumption about the fitness values thus far is that all mutants are less fit than the wild type (*w*_*i*_ < *w*_0_ = 1 for *i* ≠ 0). Then the recursions describing the dynamics of mutant frequencies over time are [38]:

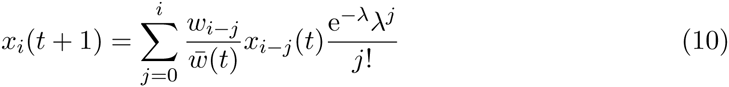

where 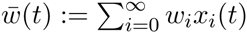 is again the population mean fitness.

Similarly to the finite locus case, we extend this model to a distribution of mutation rate under the two inheritance assumptions: If mutation rate is not inherited, then

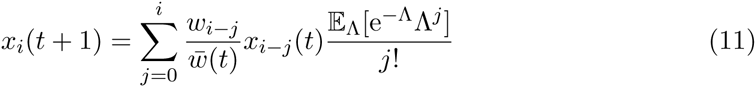

while if mutation rate is perfectly inherited, with rate *λ*_*k*_ in the *k*^th^ subpopulation at frequency *q*_*k*_, then the mutant frequencies 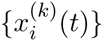 in the *k*^th^ subpopulation are given by Equation 10 with *λ* ≡ *λ*_*k*_, and the overall mutant frequencies in the population are given by

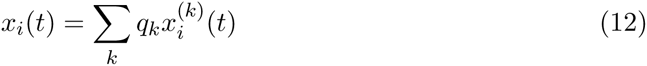

For most of our results, we will deal with a special case of the model in which *w*_*i*_ = (1 – *s*)^i^ [34], meaning that each mutation has an equal effect (cost *s*) and there is no epistasis.

## 3 Results

We are interested in the effect of mutation rate heterogeneity on the production and maintenance of mutations in a population’s standing genetic variation. We therefore focus on comparing a “heterogeneous” population, where the mutation rate (per locus, *U*, or per genome, Λ) has a given distribution, to a baseline “homogeneous” population with mutation rate fixed to the mean of this distribution (denoted 〈*U*〉 or 〈Λ〉, respectively). We first consider the probability distribution of the number of mutations occurring “simultaneously”, i.e. on a single genome in one generation, which is independent of their fitness effects. We then consider the temporal dynamics of genotypes over multiple generations of mutation and selection against deleterious mutations, and derive the mutation-selection balance attained when mutation rate varies among individuals.

### 3.1 Probability of simultaneous mutations

The probability *p*_*n*, *j*_(*u*) of *j* simultaneous mutations among *n* loci available to mutate, given a mutation rate of *u* per locus, is given by Equation 1. Averaging over the distribution of mutation rate, the overall probability of *j* simultaneous mutations is then *p*_*n*,*j*_ = E_*U*_[*p*_*n*,*j*_ (*U*)]. These probabilities can also be interpreted as the expected frequencies of *j*-point mutants produced (before selection) by a purely wild type starting population.

To consider the effect of a mutation rate that varies about its mean, we apply Jensen’s Inequality to the functions *p*_*n*,*j*_(*U*). If *g* is any real convex function of a random variable *U* (i.e. *g"*(*U*) > 0), then Jensen’s Inequality states that 〈*g*(*U*)〉 ≥ *g*(〈*U*〉), with equality if and only if *g* is linear or *U* takes on a fixed value [14, p. 27]. Thus, we can determine whether variability in mutation rate increases or decreases *p*_*n*,*j*_ ≡ 〈*p*_*n*,*j*_(U)〉, relative to *p*_*n*,*j*_(〈*U*〉) in the homogeneous case, by analyzing the second derivative of *p*_*n*,*j*_(*U*) for each *n* and *j*.

If we consider a single locus (*n* = 1), it is clear that the functions *p*_1,*j*_(*U*) are linear.Thus, the overall probability of mutation at a single locus is fully determined by the mean mutation rate and independent of the extent of variability: specifically *p*_1,0_ = 1 – 〈*U*〉 and *p*_1,1_ = 〈*U*〉. On the other hand, if we consider multiple loci (*n* ≥ 2), the functions *p*_*n*,*j*_(*U*) are non-linear, and in general 〈*p*_*n*,*j*_(*U*)〉 ≠ *p*_*n*,*j*_(〈*U*〉).

Since *p*_n,0_(*U*) = (1 – *U*)^*n*^ and *p*_*n*,*n*_(*U*) = *U*^*n*^ are clearly convex for *n* ≥ 2, we can conclude that the probabilities of either all or none of the loci mutating are increased by variability in *U*. Logically, the probability of at least some intermediate numbers of mutations must be reduced. We find (Suppl. Text II.1) that *p*_*n*,*j*_ will generally be increased by heterogeneity for the smallest and largest values of *j*, and decreased in some intermediate range of *j*, with the exact switching points depending on *n* and on the particular distribution of *U*. For realistic ranges of mutation rate in most organisms, heterogeneity will increase the chance of zero or of two or more simultaneous mutations and decrease the chance of a single mutation occurring, even with many loci under consideration. For certain RNA viruses with high mutation rates, and possibly cellular populations containing strong mutators, the switching points may be shifted upward (Figure 1).

**Figure 1:**
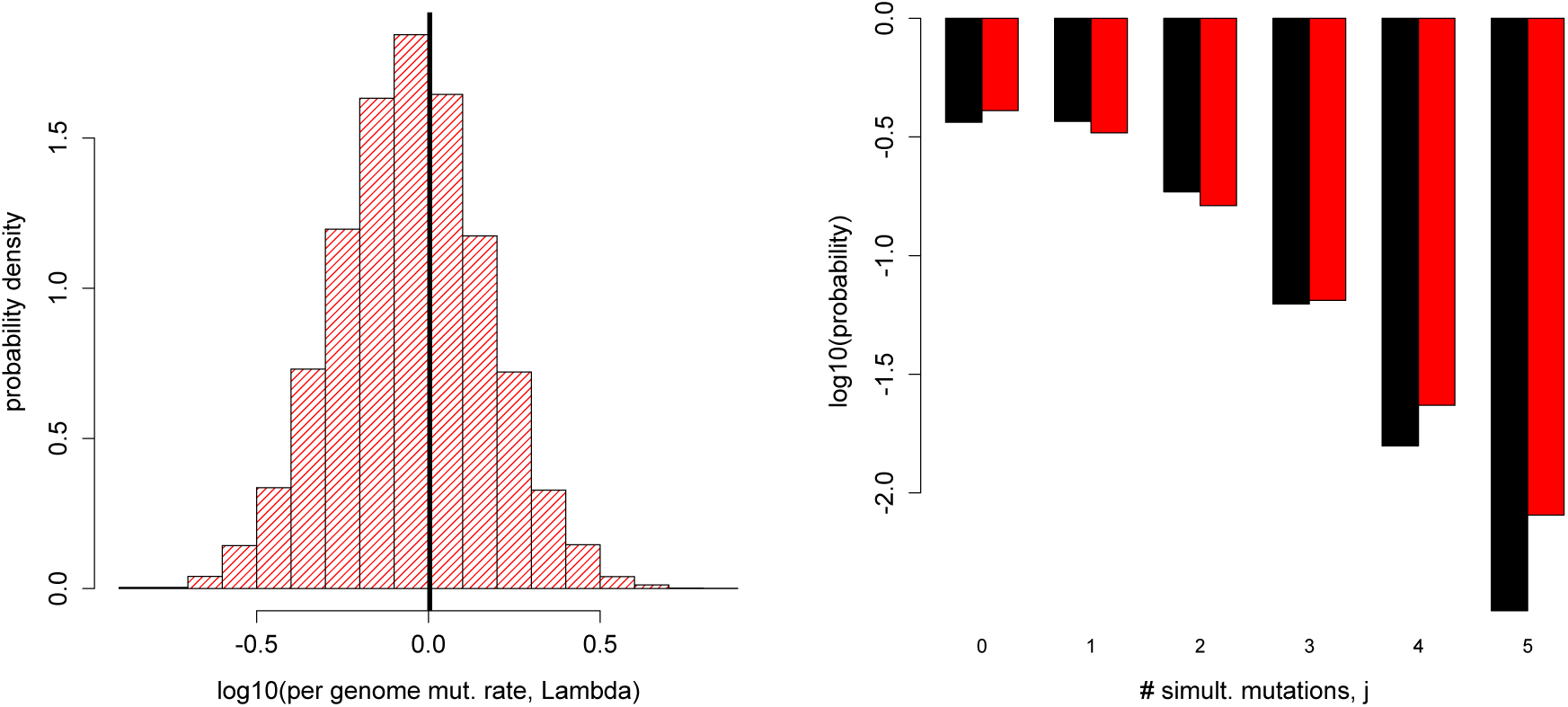
Directional effect of mutation rate heterogeneity on simultaneous mutation probability. Left: the distribution of per-genome mutation rate (Λ) obtained by sampling 10000 times from a log-normal distribution, with the thick black vertical line indicating the sample mean. Sample mean z 1:0 (similar to estimates for some RNA viruses [25]), sample variance z 0:29, range z 0:14 – 6.3. Right: probability of *j* simultaneous mutations occurring in the genome under the infinite-locus model, i.e. *p*_*j*_, if Λ is heterogeneous following the chosen distribution (red) versus fixed to the sample mean (black). The directional effects agree with those predicted analytically (keeping in mind that higher probabilities will be less negative on the log scale).

Summary statistics of the probability distribution behave as intuitively expected: the mean number of mutations occurring simultaneously on a single genome is unaffected by variability of the mutation rate about its mean, but the variance in the number of mutations is increased by variance in the mutation rate distribution (exact expressions in Suppl. Text II.1.2).

We further analyze the magnitude of the effect of heterogeneity in the case of all *n* loci mutating simultaneously. Rewriting this probability (Suppl. Text II.1.3):

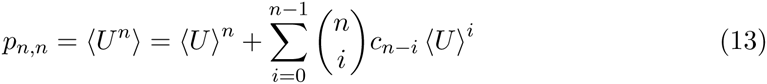

where *c*_*i*_ = 〈(U – 〈*U*〉)^*i*^ is the *i*^th^ central moment of the mutation rate distribution. (In particular, *c*_0_ = 1, *c*_1_ =0, and *c*_2_ is the variance.) Thus the probability of *n* simultaneous mutations depends on the first *n* central moments. For instance, the “boost” in triple mutations increases with the variance, and is larger when the distribution is right-skewed than when it is left-skewed. Note that even-numbered central moments must be positive, while odd-numbered central moments may be positive or negative; however, according to Jensen’s Inequality, any negative terms must be outweighed by the positive terms. It can also be shown (Suppl. Text II.1.3) that the *relative* effect of heterogeneity in mutation rate on the probability of simultaneous mutation at all loci increases with the number of loci under consideration (Fig. 2).

**Figure 2:**
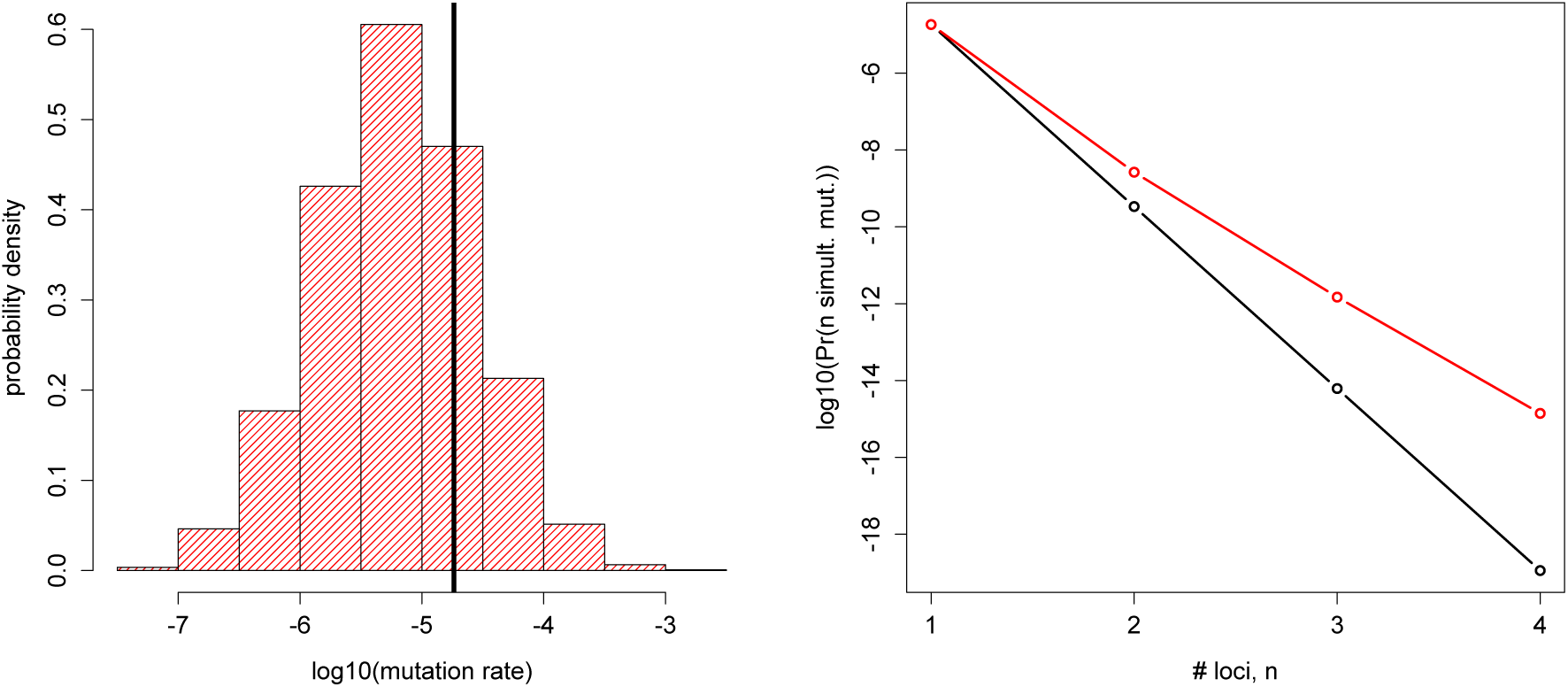
Relative effect of mutation rate heterogeneity increases with the number of mutating loci. Left: the distribution of per-locus mutation rate (*U*) obtained by sampling 10000 times from a log-normal distribution, with the thick black vertical line indicating the sample mean. Sample mean z 1:8 × 10^‒5^ (close to the base substitution rate estimated for HIV [54]), sample variance z 2:3 × 10^‒9^, range z 3:2 × 10^‒8^ – 1:4 × 10^‒3^. Right: probability of all *n* loci under consideration mutating simultaneously, i.e. *p*_*n,n,*_ as a function of *n*, if *U* is heterogeneous following the chosen distribution (red) versus fixed to the sample mean (black).

### 3.2 Deterministic mutant frequency dynamics under mutation and selection

We now consider genotype frequencies over multiple generations of mutation and selection, with particular attention to the equilibrium (mutation-selection balance). This deterministic approach neglects demographic stochasticity in a finite population, and importantly in our extension, also neglects sampling effects from the mutation rate distribution. We compare the well-known results for fixed mutation rate to our novel results for heterogeneous mutation rate. Details of the mathematical results are provided in Supplementary Text II.2.

#### 3.2.1 One locus

Heterogeneity in the mutation rate again turns out to have a negligible effect on mutant frequency dynamics at a single focal locus. In particular, the classic mutation-selection balance is simply replaced by 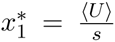 where 〈*U*〉 is the mean mutation rate in the population and s is the cost of the mutation. Population mean fitness at equilibrium is correspondingly given by *w̄*^*^ = 1 – 〈*U*〉. These solutions are valid regardless of whether mutation rate is inherited.

The full temporal dynamics of the mutant frequency are more involved, but still amenable to analytical solution. In the non-inherited case, the temporal solution exactly coincides with the homogeneous case. Mathematically, this is because the relevant mutation-selection matrix is identical: due to the linearity of *M*(*U*) in *U* (Equation 5), we have 〈*M*(*U*)〉 = *M*(〈*U*〉). In the inherited case, the solutions are not exactly equivalent, but can be shown to coincide up to first order in the maximum mutation rate. Mathematically, the mutant frequency in each subpopulation at time *t* is nonlinear in *u*_*k*_, and this nonlinear expression must be averaged over the distribution of mutation rate; however, higher order terms make a negligible contribution (vanishing at equilibrium).

#### 3.2.2 Multiple loci

When multiple loci are involved, the mutation-selection matrix *M* becomes non-linear in *U*. Specifically, in the *n*-locus model terms up to order *U*^*n*^ appear in M, and we expect the highest-order mutants to have frequency with leading order *U*^*n*^. Thus, the first *n* moments of the mutation rate distribution will generally play a non-negligible role in genotype frequency dynamics. We conduct a detailed mathematical analysis of the two-locus case, while a brief consideration of the infinite-locus limit confirms our key qualitative conclusions and suggests how results will extend to more loci. Below we focus on key results and their intuitive interpretation, while detailed expressions for genotype frequencies over time in all model cases are provided in Supplementary Text II.2. Throughout, *V* denotes the variance of the mutation rate distribution and *c*^2^:= V/ 〈*U*〉^2^ denotes the squared coefficient of variation. To distinguish models under comparison, the short form ‘het’ will indicate a heterogeneous mutation rate characterized by a distribution, and ‘hom’ a homogeneous mutation rate fixed to the mean of this distribution. Further, *H* = 0 will indicate that mutation rate is non-inherited and *H* = 1 will indicate perfect inheritance.

##### Genotype frequencies in the two-locus model

For reference, with fixed mutation rate *U* ≡ *u*, the equilibrium frequencies are approximated to order *u*^2^ by:

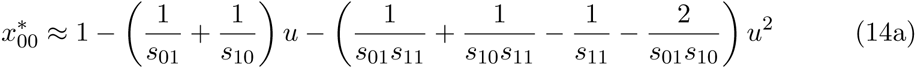

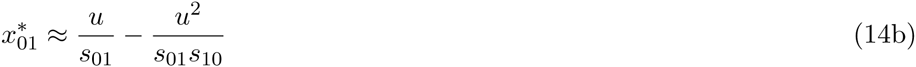

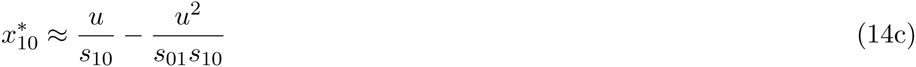

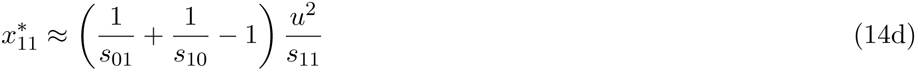

and the population mean relative fitness is given by *w*̄^*^ = (1 – *u*)^2^.

When heterogeneity is introduced, we use the recursions given in Equations 7 (no inheritance) or 8-9 (perfect inheritance) to solve for the genotype frequencies. Figure 3 illustrates examples of the resulting temporal dynamics of double mutant frequency, *x*_11_ (*t*). Our analytical approximations (Suppl. Text II.2.2) typically show excellent agreement to results obtained by numerical iteration of the recursions. In all cases, the double mutant frequency is elevated by variability of the mutation rate about the mean. This increase is larger (after the first generation) when mutation rate is perfectly inherited. However, while the non-inherited case falls substantially short for weak selection against single mutants, the two cases become similar for strong selection against single mutants. Our analytical solutions indicate that these observations are general: double mutant frequency is increased by an amount proportional to variance in mutation rate, and depends on the selection coefficients in a way that will be clarified below. The equilibrium solutions are summarized in Table 1 for reference. As a note of caution, the error in these approximations scales with max(〈*U*〉^3^, 〈*U*〉 *V*, *V*^2^) in the case of non-inheritance and with 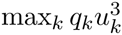 in the case of perfect inheritance. Further numerical testing (not shown) indicates that the approximations can break down for extreme mutation rate distributions, particularly when at least one selection coefficient is small.

**Figure 3:**
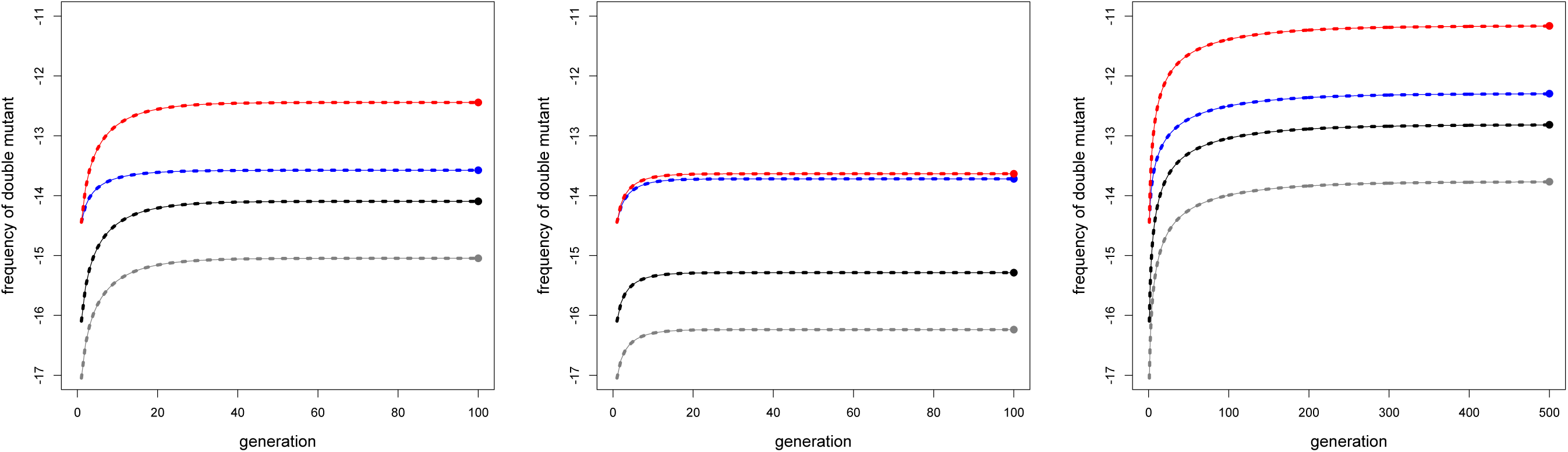
Temporal dynamics of the double mutant with approach to equilibrium, assuming the population is initially composed entirely of the wild type. Mutation rate takes on either of two values: *U* = *u*_*l*_= 3 × 10^‒9^ with probability 0.99 or *U* = *u*_*h*_ = 6 × 10^‒7^ with probability 0.01, thus 〈*U*〉 z 9.0 × 10^‒9^ and *V* z 3.5 × 10^‒15^. These values are reasonable for loci with large target sizes in a bacterial population containing a strong hypermutator at 1% frequency, parameterizing approximately from [43]: *u*_*l*_ is the wild type *E. coli* mutation rate to rifampicin resistance estimated from a uctuation assay and *u*_*h*_=*u*_*l*_ is the fold-increase in mutation rate in the mismatch repair defective MutL^‒^strain. The selection coeffcients vary across panels: left, *s*^01^ = *s*^10^ = 0:1 and *s*^11^= 0:19; center, *s*^01^ = *s*^10^ = 0:9 and *s*^11^ = 0:19; right, *s*^01^ = *s*^10^ = 0:1 and *s*^11^ = 0:01. Black indicates the homogeneous case with mutation rate fixed to 〈*U*〉; blue is the heterogeneous case with no inheritance; and red is the heterogeneous case with perfect inheritance. Grey additionally shows the result when *U*≡*u*_l_. Thus comparing black to grey indicates the effect of changing the mean mutation rate by adding a hypermutator, while comparing blue/red to black isolates the effect of increasing variance with fixed mean. The solid line in each case indicates the result of numerically iterating the recursions, while the dashed line indicates the analytical approximation for the temporal dynamics, and the large point at the end of each curve indicates the analytical approximation of the equilibrium.

**Table 1:**
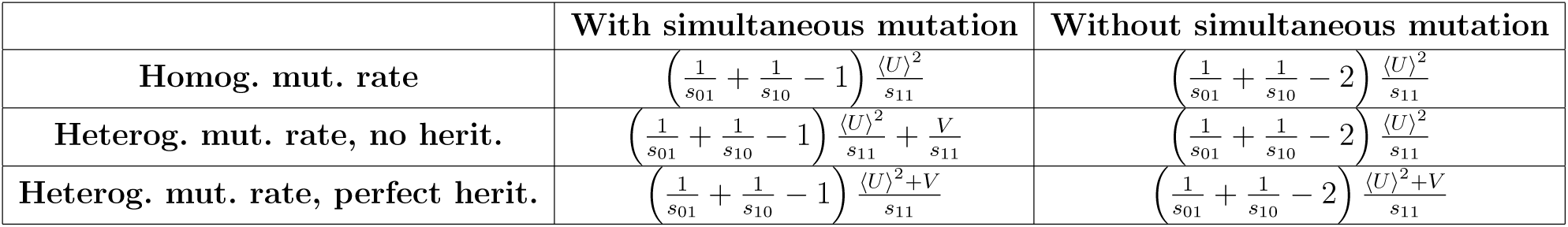
Approximate equilibrium frequency of double mutants 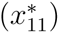 in various cases of the two-locus model. In the homogeneous case, we suppose the mutation rate *u* is fixed to 〈*U*〉 for comparison with the heterogeneous cases, where the equilibrium is defined by both the mean 〈*U*〉 and variance *V* of the mutation rate distribution.

Heterogeneity also affects the frequencies of the wild type and single-point mutants. Generally, heterogeneity in the mutation rate has the effect of clustering mutations at mutation-selection balance, i.e. increasing the frequency of the double mutant at the expense of single mutants, similar to the qualitative effect on the production of new mutations in each generation. More specifically, if mutation rate is not inherited, heterogeneity decreases the equilibrium frequency of single mutants and increases that of the wild type. If mutation rate is perfectly inherited, heterogeneity still decreases the equilibrium frequency of single mutants, but interestingly, the effect on the wild type can take either direction. Namely, 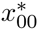 is decreased when epistasis is sufficiently strongly positive, in which case the additional double mutants appear to exert sufficient competition on the wild type to outweigh the increased chance of mutation-free reproduction in each generation.

##### Understanding the roles of mutation rate inheritance and simultaneous mutations

Our results can be understood by considering which of the underlying mutational pathways to the double mutant are affected by heterogeneity in the mutation rate. In the absence of inheritance, the mutation rate experienced by multiple loci on a genome only shows an association over one generation. Any boost in multiple mutant frequency due to mutation rate heterogeneity must thus be achieved through a boost in simultaneous mutations. In contrast, effects of heterogeneity can act across generations when the mutation rate is inherited. Then multiple mutants can be boosted not only by simultaneous mutations, but also by stepwise accumulation of mutations over several generations.

To confirm this reasoning we consider a variant of the mutation model, where simultaneous double mutations are not allowed. Then higher-order terms in *U* no longer appear in the mutation-selection matrix (Equation S10 in Suppl. Text II.2.2.5), indicating that in the absence of inheritance, only the mean mutation rate matters. This is in line with our insight that all effects of non-inherited variation in mutation rate must act via the eliminated simultaneous mutations. On the other hand, when mutation rate is perfectly inherited, variance can still have an effect, albeit reduced, via stepwise accumulation of mutations. Mathematically, multiplication of the mutation-selection matrix (i.e. iteration over generations) gives rise to higher-order terms in *U* before the result is averaged over the mutation rate distribution.

We are now in a position to interpret the increases in double mutant frequency due to mutation rate heterogeneity. We denote the absolute increase by

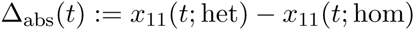

and the relative increase by

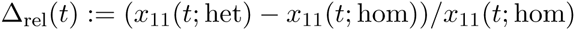

It turns out that Δ_abs_(*t*) is proportional to the variance (*V*) and Δ_rel_(*t*) is proportional to the squared coefficient of variation (*c*^2^ = *V*/ 〈*U*〉^2^) at all times *t* (Suppl. Text II.2.3.1).

With perfect inheritance of mutation rates, mutation rate heterogeneity affects all pathways equally. Thus selection plays the same role as in the homogeneous case and the double mutant frequency is simply scaled up by a constant factor:

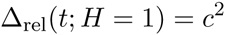

independent of *t* and {*s*_*i*_}. Plotting the equilibrium frequency 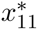 as a function of single mutant cost, this effect manifests as parallel curves in the homogeneous and heterogeneous cases (Fig. 4). When single mutants are sufficiently fit, stepwise accumulation of mutations is the main pathway, and blocking simultaneous mutations has little effect on the double mutant frequency; however, when single mutants are very costly, blocking simultaneous mutations has a drastic effect. The precise ranking of 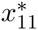 all model cases, as determined by {*s*_*i*_} and *c*^2^, is given in Supplementary Text II.2.3.2. Importantly, since the perfectly inherited case yields the maximal increase compared to the homogeneous case, we can conclude that *c*^2^ in fact provides an upper bound on the relative increase in double mutant frequency that can be achieved by mutation rate heterogeneity across all choices of selection coefficients, times (since starting from a wild type population), and inheritance assumptions.

**Figure 4:**
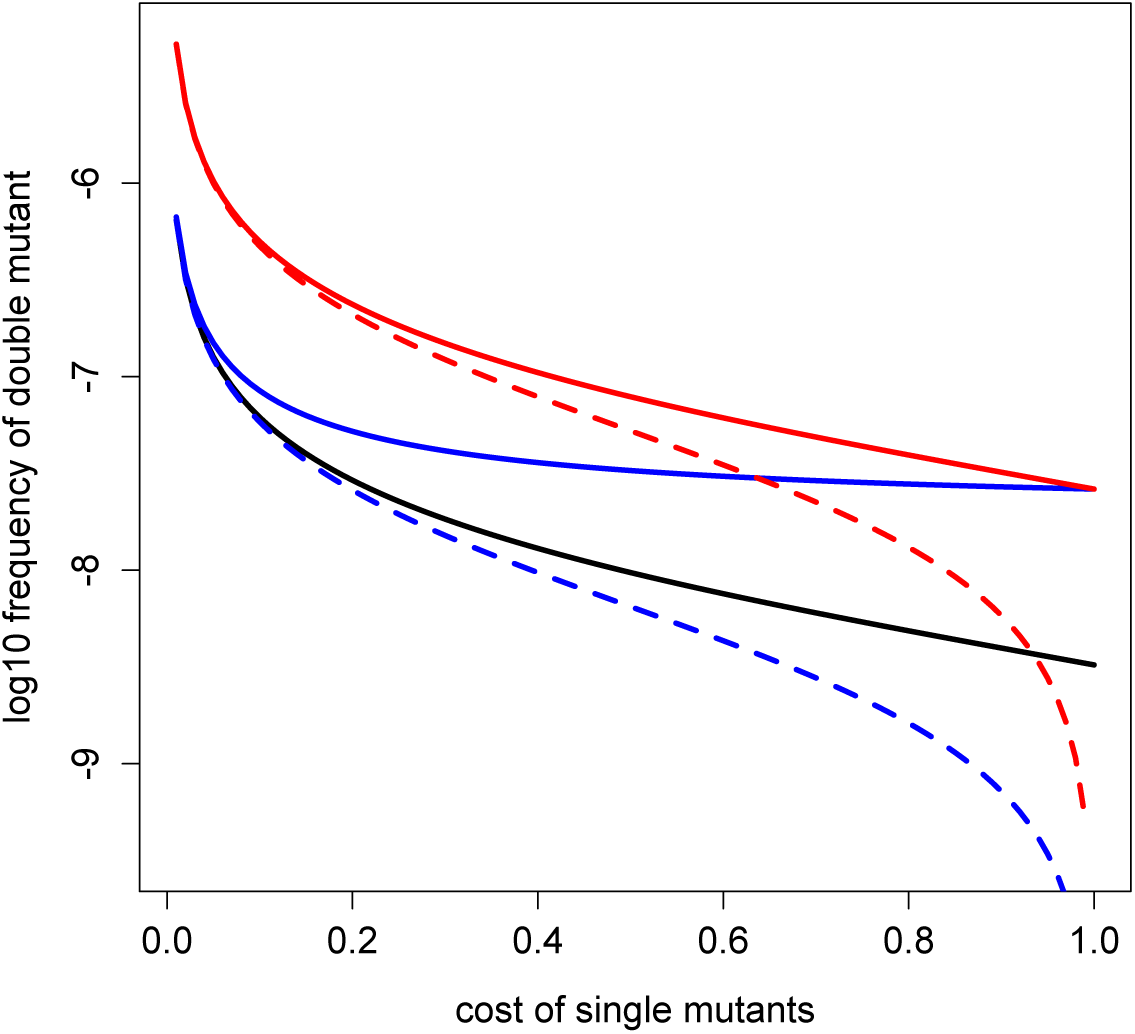
Equilibrium frequency of the double mutant as a function of single mutant costs. The analytical approximations for the equilibrium double mutant frequency (Table 1)are plotted for the various model cases: homogeneous – black; heterogeneous, no inheritance–blue; heterogeneous, perfect inheritance – red. Solid lines indicate the result when simultaneous mutations are allowed; dashed lines when simultaneous mutations are blocked. We take 〈*U*〉 = 1.8 × 10^‒5^ and *V* = 2.3 × 10^‒9^ as in Figure 2.

In the non-inherited case, the precise increase in double mutant frequency depends on the selection coefficients. (We focus here on the equilibrium; transient dynamics are analyzed in Suppl. Text II.2.3.1.) The absolute increase

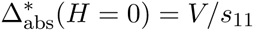

is due entirely to the increased influx of simultaneous double mutations, which are filtered by selection according to coefficient *s*^11^. The selection coefficients of the single mutants do not appear, because stepwise accumulation of mutations cannot be affected by variance in mutation rate in this case. However, the relative increase

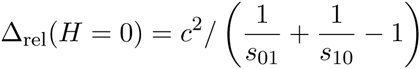

increases with *s*_01_ and *s*_10_, because the stronger the selection against single mutants, the fewer the double mutants can be produced via these intermediates, and the more relatively important simultaneous double mutation becomes. In Figure 4, the heterogeneous noninherited case thus approaches the homogeneous case as *s*_01_ = *s*_10_ → 0, and approaches the heterogeneous perfectly inherited case as *s*_01_ = *s*_10_ → 1, where all double mutants must be generated directly by simultaneous mutation from the wild type. When simultaneous mutation is blocked, the heterogeneous non-inherited case drops to align perfectly with the homogeneous case over the full range of single mutant costs.

##### Genotype frequencies in the infinite-locus model

The two-locus results are supported and extended by considering an infinite-locus model. For simplicity, we consider the special case in which each individual mutation has an equal fitness effect (cost *s*) and there is no epistasis. In this case, for fixed per-genome mutation rate *λ*, the equilibrium frequency of *i*-point mutants is given by [34]:

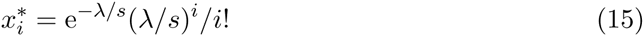

The equilibrium frequencies thus take the same form as the probabilities of simultaneous mutation, namely a Poisson distribution, but now with parameter *λ*/*s* instead of *λ*.

With a heterogeneous mutation rate, in the case of perfect inheritance, one must simply average the fixed-rate solution over the distribution of Λ. Thus an analysis of the effect of heterogeneity, applying Jensen’s Inequality, proceeds exactly as in the simultaneous mutation analysis: Heterogeneity in mutation rate has the effect of increasing the frequency of zero‐ or possibly few-point mutants, decreasing the frequency of intermediate mutants, and increasing the frequency of many-point mutants. Moreover, decreasing s has the same effect as increasing the mutation rate, namely shifting upwards the values of *i* at which the directional effect of heterogeneity switches sign. In the case of non-inherited mutation rate, analytical progress does not appear straightforward, but we intuitively expect the same qualitative effect. We confirm this pattern by numerical iteration of the recursions for both cases (Equations 10-12). The switching points in the directional effect appear as predicted in both cases, but are shifted upward when mutation rate is noninherited (Fig. 5). Furthermore, among the higher mutant classes where heterogeneity boosts mutant frequency, the relative magnitude of this effect increases with number of mutations, as also seen in the simultaneous mutation results.

**Figure 5:**
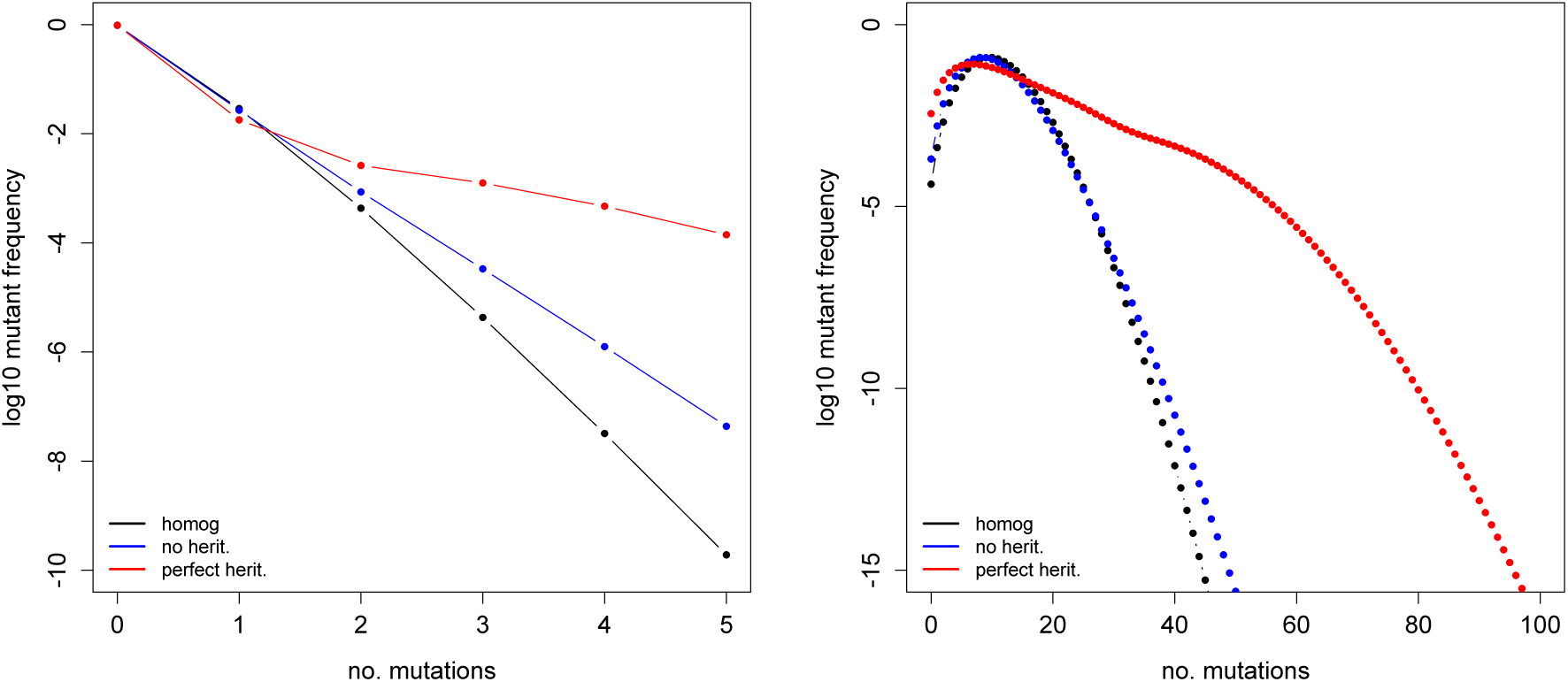
Equilibrium mutant frequencies under the infinite-locus model. Specifically, the base 10 log of the genotype frequency at the numerically determined equilibrium is plotted as a function of number of mutations carried, in each model case (homogeneous – black; heterogeneous with no inheritance – blue, or perfect inheritance – red). The selection coeffcient per mutation is *s* = 0:1. The results are illustrated for two example mutation rate distributions. Left: Λ takes on two values, 0.0015 with probability 0.99 or 0.15 with probability 0.01 (mean ~ 0:003 is bacteria-like; [24]). Right: Λ is given by 1000 draws from a log-normal distribution. Sample mean is 1.01 (RNA‐ or retrovirus-like; [24]), sample variance is 0.29, and range is 0.19-4.2.

Thus, in qualitative agreement with the two-locus results, heterogeneity has the effect of clustering mutations at mutation-selection balance. Furthermore, heterogeneity again appears to have a larger impact on the mutant frequency distribution when the mutation rate is inherited. More specifically, there is a larger increase in the frequency of the lowest‐ and highest-order mutants (and correspondingly larger decrease in the frequency of intermediates). Again, the relative importance of inheritance varies with the strength of selection: the non-inherited case appears more similar to the homogeneous case when *s* is smaller, and gradually becomes more similar to the perfectly-inherited case as *s* increases (Fig. S1).

Clearly, increasing the variance of the mutation rate distribution generally increases the effects of heterogeneity (Fig. S1). However, the differences in mutant frequencies are no longer directly proportional to variance, since all higher moments of the mutation rate distribution also affect the results in the infinite-locus case.

##### Frequency of mutant alleles and average number of mutations per genome

We have seen that heterogeneity in mutation rate clusters mutations at multiple loci. We now ask whether it has any effect on the frequency of the mutant allele at any given locus 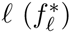 or on the average number of mutations carried per genome 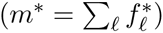 at equilibrium.

In the two-locus model, it turns out that both these quantities differ between the heterogeneous and homogeneous cases by an amount proportional to the variance in mutation rate. However, the magnitude and direction of the effect depends on epistasis and mutation rate inheritance (details in Suppl. Text II.2.2.4). With perfect inheritance, variance increases the frequency of any given mutant allele if and only if epistasis (ϵ) is positive. Intuitively, since variance in the mutation rate increases double mutant frequency at the expense of single mutants, the overall frequency of the mutant allele at any individual locus will be elevated precisely when the double mutant is fitter than expected from the singles. Without inheritance, the frequency of the mutant allele is increased by variance when the double mutant is fitter than the single mutant, which can be translated into a more stringent condition on epistasis than in the perfect-inheritance case (ϵ > ϵ_*c*_ > 0). Similar effects arise for *m*^*^.

In the infinite-locus model, every mutation occurs at a unique site, so it no longer makes sense to consider 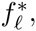, but we can still consider *m*^*^. We analyze only the special case of the model where every mutation has equal cost and there is no epistasis. Then consistent with the two-locus finding, in the case of perfect inheritance *m*^*^ is not affected by heterogeneity in the mutation rate (Suppl. Text II.2.4.2). In the case of no inheritance, extrapolating from the two-locus model, we expect that *m*^*^ is reduced by heterogeneity in the absence of epistasis. Although we lack an analytical result, this indeed appears numerically to be the case (confirmed for the cases illustrated in Figures 5 and S1).

##### Mutational load

In the two-locus model, the population mean fitness at equilibrium is given by:

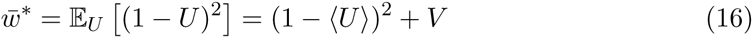

regardless of whether mutation rate is inherited (Suppl. Text II.2.2). Thus, mutational load is decreased by an amount equal to the variance of mutation rate in the population. This effect can be explained by the clustering of mutations into fewer individuals.

In the limiting infinite-locus model in which fitness is determined by number of mutations (with otherwise arbitrary fitness costs), the population mean fitness at equilibrium is given by

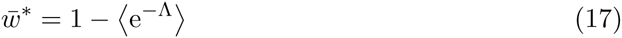

again regardless of inheritance. An application of Jensen’s Inequality demonstrates that *w̄*^*^ is likewise enhanced, i.e. mutational load is reduced, by variability in the mutation rate.

## 4 Discussion

### 4.1 Significance

The critical role of mutations in producing the raw material for evolution has long been recognized by biologists and mathematically analyzed by population geneticists. In particular, the maintenance of standing genetic variation is known to be shaped by both mutation rate and selection coefficients. While the distribution of mutational fitness effects has been intensively studied, the distribution of mutation rates among members of a population has received far less attention. Heterogeneity in mutation rate could stem from a wide range of sources, including genetic differences, environmental influences, and random physiological fluctuations. In this study, we showed that such heterogeneity has potentially far-reaching evolutionary consequences by affecting the chance of multi-point mutants appearing, the frequency of mutant genotypes harbored in the standing genetic variation, and the population mean fitness. Our general modeling approach allowed an arbitrary distribution of mutation rate, clearly separated the role of variation about the mean from effects on the mean itself, and compared inherited versus non-inherited forms of variation. Our first key finding is that mutation rate heterogeneity results in an overrepresentation of higher-order mutants and an under-representation of intermediate mutants. We gained a quantitative understanding of these effects as a function of moments of the mutation rate distribution and selection coefficients of the mutants. Our second key finding is that, due to this clustering of mutations into fewer individuals, the population as a whole has a reduced mutational load.

The qualitative effect that variability in mutation rate among individuals should result in an over-representation of multiple-point mutants has previously been recognized [62, 23, 22, 27], but our rigorous analysis of the quantitative conditions for this effect to occur is novel. Moreover, while previous calculations of the contribution of mutators have implicitly only considered simultaneous mutations [62, 11, 23], we also analyzed the effects on accumulated mutant frequency when both *de novo* mutation and selection act over time. We thus showed that heterogeneity in mutation rate increases not only the probability of multiple mutations arising simultaneously on a genome (Section 3.1), but also the frequency of multiple mutants at mutation-selection balance (Section 3.2). In Supplementary Text III, we additionally show numerically that heterogeneity reduces the waiting time for the first appearance of double mutants in a stochastic branching process model.

Interestingly, this skewing of the mutant frequencies leads to the appearance of positive linkage disequilibrium. If detected empirically, the standard interpretation of such an observation would be the existence of epistasis. However, our results show that an alternative explanation is the existence of population heterogeneity in mutation rate. Drake and colleagues observed that multiple mutants often appear to be over-represented (relative to a Poisson distribution) in mutant counts from experiments, and indeed interpreted this finding as a sign of heterogeneity in the mutation rate [23, 22]. However, it was unclear whether selection could be ruled out in all the reported experiments, and thus whether epistatic fitness effects might also have boosted double mutant frequency. An interesting direction for future work would be to disentangle the effects of mutation rate heterogeneity and epistasis, with a rigorous statistical analysis of the existing experimental findings.

We derived expressions for mutant frequency that both indicate the key parameters at play and allow a quantitative estimation of their effects. The dynamics of mutants at *n* focal loci are driven by the first *n* moments of the mutation rate distribution. Thus if we examine only one locus, the population mean mutation rate is sufficient to predict mutant dynamics, but to predict the joint dynamics at two loci, variance must be considered, and so on. Specifically, for two loci, we found that the frequency of double mutants is boosted (compared to a population with mutation rate fixed to the same mean) by an absolute amount proportional to the variance, or a relative amount proportional to (and at most equal to) the squared coefficient of variation. Variance likewise appeared to determine the reduction in waiting time for double mutants in the stochastic model (Suppl. Text III.2 and Fig. S4). The quantitative effects can be substantial. For example, parameterizing from data on mutation rates to rifampicin resistance in bacteria [43], the presence of hypermutator individuals with 200-fold increased mutation rate at 1% frequency is predicted to increase the equilibrium frequency of double mutants by ~9-fold through effects on the mean, but up to ~45-fold further through effects on the variance at fixed mean (Fig. 3 and analytical solutions in Section 3.2). On the other hand, more evenly distributed mutation rates result in smaller effects; for instance, using the best-fitting per base pair mutation rate distribution inferred in a population of budding yeast [37], variance increases double mutant frequency by maximally 1.4-fold relative to a homogeneous population with the mean mutation rate.

Moreover, we predict that mutation rate heterogeneity becomes increasingly significant with more loci under consideration, in that the relative boost in frequency is larger for higher-order mutants. This suggests that “mutators” in a population could make a much greater contribution to adaptation than suggested by analyses focusing on single loci (see also [62]). Furthermore, one should proceed with caution in applying mutation rate estimates obtained by assaying a phenotype that can be conferred by a single point mutation. If mutation rate is heterogeneous in the population, such an assay will in fact estimate the mean rate, a reasonable result in itself. However a naive extrapolation assuming this rate to be fixed would underestimate the chance that such a population will harbor multiple mutants. This is particularly concerning for analyses of whether multidrug resistant pathogens or cancerous cells are likely to “pre-exist” before a patient starts a drug treatment [67, 41, 12]. Progression to cancer via accumulation of multiple mutations may also occur faster than expected in a cellular population, even when controlling for changes in mean mutation rate. (The importance of generally elevating mutation rate in cancerous cells has previously been pointed out; [49, 46, 47, 48].)

Our analysis also clarified the role of parent-offspring correlation of mutation rate, by directly comparing cases where mutation rate is either perfectly inherited or drawn independently at random by each individual, such that the population-level distribution is the same in both cases. (Realistically, different underlying mechanisms are likely to give rise to both different levels of correlation and different distributions of mutation rate, but we construct this hypothetical scenario to isolate the effect of the former.) Realistically, parent-offspring correlation falls on a broad spectrum between the two extremes analyzed here, and thus the effects of mutation rate variation are likely to be intermediate. The more strongly mutation rate is correlated through a lineage, the greater the effect of variation about the population mean, via the boosted rate of stepwise accumulation of mutations. When mutation rates are uncorrelated, variation about the mean can only have an effect through increasing the chance of simultaneous mutations, since non-random association of mutation rate at multiple loci on a genome only lasts for one generation. A previous study concluded that simultaneous mutations make a negligible contribution to multilocus adaptation [50], but their results supposed that single mutants are neutral or only slightly deleterious. We considered a wider range of effects and found that if intermediates are highly deleterious, simultaneous mutations play a crucial role in generating multiple mutants directly. In this case, the extent to which mutation rate is inherited makes little difference, so long as the population as a whole contains individuals with elevated mutation rate (Section 3.2 and Suppl. Text III). The case of low-fitness intermediates would be highly relevant for instance when a multi-drug combination therapy is applied to a pathogen population, and each mutation confers resistance to a single drug but no protection against the other drugs.

Taken together, our results suggest that populations with heterogeneous mutation rate have a greater capacity to adapt. Firstly, they generate multi-point mutants faster, which applies to beneficial *de novo* mutations as well as deleterious ones. In particular, crossing fitness valleys is expected to be accelerated (see also [2, 50]). Secondly, in the long term they harbor costly multi-point mutants at a higher frequency in the standing genetic variation. This diversity is expected to be an important source for adaptation if the environment changes and previously deleterious genotypes become favorable [5].

Finally, the clustering of multiple mutations into fewer individuals implies that a population with heterogeneous mutation rate not only can explore genotype space more widely, but simultaneously maintains higher population mean fitness. Such a possibility has previously been suggested verbally [23, 13] and uncovered in particular models assuming mutation rate can take on two possible values, with no parent-offspring correlation [29, 2]. We find that this effect is much more general, holding for any distribution of mutation rate, and, perhaps surprisingly, independently of the degree of parent-offspring correlation. Furthermore, it does not require that mutation rate is specifically elevated in individuals with low fitness, which was previously found to yield a similar effect of simultaneous “adaptedness and adaptability” in a model of stress-induced mutagenesis [66]. Thus, besides concentrating high mutation rates into limited time periods or limited parts of the genome [72], concentrating high mutation rates into particular individuals in a population (whether heritably or not) adds another possible solution to the conundrum of balancing gains in adaptability against fitness losses through deleterious mutations

### 4.2 Model limitations and extensions

We point out a few noteworthy assumptions of our modeling approach. Firstly, we described an asexually reproducing haploid population. We expect similar but weaker qualitative effects to hold in sexual populations. Recombination can bring together mutations generated in different lineages, thus reducing the importance of rare hypermutators in accelerating the first appearance of multi-point mutants. In the longer term, recombination would counteract positive linkage disequilibrium and thereby reduce the excess multiple mutants generated by mutation rate heterogeneity.

Secondly, we neglected any fitness effects directly associated with mutation rate, including any dependence of mutation rate on the alleles carried at the focal loci. However, particularly for viruses, there is a correlation between replicative fitness and mutation rate, since both are influenced by the speed of replication of the genetic material [28, 15]. Furthermore, environmental heterogeneities that may affect mutation rate (such as chemotherapeutic drugs) also are likely to affect fitness. The level of “stress” experienced by an individual and hence its mutation rate could then depend on its genotype: for example, bacteria appear to down-regulate the SOS stress response (associated with mutagenesis) as they adapt to a stressful environment [77].

Finally, we assumed that the mutation rate distribution was fixed from generation to generation. Nonetheless, temporal environmental changes or evolution of the mutation rate could yield changes in the population’s distribution of mutation rate over time. Analyzing the joint effects of inter-individual variation and population-level temporal changes in mutation rate would be an interesting direction for future work.

### 4.3 Empirical Outlook

Given the evolutionary significance we predict for mutation rate heterogeneity, it would clearly be of interest to gain a better empirical understanding of this heterogeneity. There are two approaches to this question. Firstly, mutation rate as a function of some genetic or environmental variable can be quantified using standard techniques to estimate (mean) mutation rate in a population under each fixed condition. Combining this functional relationship with a measurement or model of how the relevant variable is distributed in a natural population would suggest in turn how mutation rate is distributed. The latter point could be achieved for instance by expanding on recent advances in isolating single virus-infected host cells [13] or single cells from cancerous tumors [79, 4, 60] and quantifying mutation rate in their descendant lineages.

While it is well established that various alleles and environmental factors can affect mutation rate, there is still work to be done to quantify these relationships in a way that is useful for parameterizing population genetic models and thereby predicting evolutionary consequences of mutation rate variation. We echo other authors [69, 51] in emphasizing the need to infer mutation rate per generation, as opposed to reporting only the mean frequency of mutants counted at the end of culture growth. While the former is a more informative and reproducible measure [69], the latter remains in frequent use in experimental studies of stress-induced mutagenesis and characterizations of natural isolates.

The second and more challenging direction is to quantify how mutation rate varies from individual to individual within a population even under fixed experimental conditions, including the detection of non-inherited variation. This requires statistical analysis of the distribution of number of mutations per individual to test for deviations from the expectation under a model with uniform mutation rate [23, 22, 27, 37]. Recently developed individual-based methods provide new ways to count mutations, including fluorescent labeling of nascent mutant foci in *E. coli* [27] and isolation and whole-genome sequencing of mother and daughter cells in budding yeast [37]. Nonetheless, an inherent challenge is that mutations are rare events, and at least double mutants are required to detect non-uniformity. Besides examining the whole genome or using mutator strains [27, 37], technical advances could make it feasible to examine more individuals in a population. In parallel, a statistical power analysis (determining the sample size required to detect mutation rate differences of a given magnitude) would be valuable both for experimental design and for retrospective interpretation of existing results.

## Acknowledgements

SB acknowledges funding by the ETH Zurich, by the European Research Council under the 7th Framework Programme of the European Commission (PBDR:Grant Agreement Number 268540), and by the Swiss National Science Foundation (Grant Nr. 155866). The authors thank members of the Theoretical Biology group at ETH Zurich for insightful comments on this work, particularly Dominique Cadosch and Antoine Frenoy for discussion of stress-induced mutagenesis, and Antoine Frenoy for comments on a draft of this manuscript.

## References

[1] B. Al-Lazikani, U. Banerji, and P. Workman. Combinatorial drug therapy for cancer in the post-genomic era. Nature Biotechnology, 30:1–13, 2012.

[2] K. Aoki and M. Furusawa. Promotion of evolution by intracellular coexistence of mutator and normal DNA polymerases. Journal of Theoretical Biology, 209:213–222, 2001.

[3] M.-R. Baquero, A. I. Nilsson, M. C. Turrientes, D. Sandvang, J. C. Galán, J. L. Martínez, N. Frimodt-Møller, F. Baquero, and D. I. Andersson. Polymorphic mutation frequencies in Escherichia coli: emergence of weak mutators in clinical isolates. Journal of Bacteriology, 186:5538–5542, 2004.

[4] L. J. Barber, M. N. Davies, and M. Gerlinger. Dissecting cancer evolution at the macro-heterogeneity and micro-heterogeneity scale. Current Opinion in Genetics & Development, 30:1–6, 2015.

[5] R. D.H. Barrett and D. Schluter. Adaptation from standing genetic variation. Trends in Ecology and Evolution, 23:38–44, 2008.

[6] I. Bjedov, O. Tenaillon, B. Gérard, V. Souza, E. Denamur, M. Radman, F. Taddei, and I. Matic. Stress-induced mutagenesis in bacteria. Science, 300:1404–1409, 2003.

[7] B. Björkholm, M. Sjölund, P. G. Falk, O. G. Berg, L. Engstrand, and D. I. Andersson. Mutation frequency and biological cost of antibiotic resistance in Helicobacter pylori. PNAS, 98:14607–14612, 2001.

[8] L. Boe. Translational errors as the cause of mutations in Escherichia coli. Molecular & General Genetics, 231:469–471, 1992.

[9] L. Boe, M. Danielsen, S. Knudsen, J. B. Petersen, J. Maymann, and P. R. Jensen. The frequency of mutators in populations of Escherichia coli. Mutation Research, 448:47–55, 2000.

[10] R. Bürger. The Mathematical Theory of Selection, Recombination and Mutation. Wiley, 1st edition, 2000.

[11] J. Cairns. Mutation and cancer: the antecedents to our studies of adaptive mutation. Genetics, 148:1433–1440, 1998.

[12] C. Colijn, T. Cohen, A. Ganesh, and M. Murray. Spontaneous emergence of multiple drug resistance in tuberculosis before and during therapy. PLoS One, 6:e18327, 2011.

[13] M. Combe, R. Garijo, R. Geller, J. M. Cuevas, and R. Sanjuán. Single-cell analysis of RNA virus infection identifies multiple genetically diverse viral genomes within single infectious units. Cell Host & Microbe, 18:424–432, 2015.

[14] T. M. Cover and J. A. Thomas. Elements of Information Theory. John Wiley & Sons, Inc., Hoboken, NJ, 2nd edition, 2006. Accessed online.

[15] M. J. Dapp, R. H. Heineman, and L. M. Mansky. Interrelationship between HIV-1 fitness and mutation rate. Journal of Molecular Biology, 425:41–53, 2013.

[16] T. Day. Modelling the ecological context of evolutionary change: déjá vu or something new? In K. Cuddington and B. E. Beisner, editors, Ecological Paradigms Lost: Routes of Theory Change, chapter 13, pages 273–309. Academic Press (Elsevier), Burlington, MA, USA, 2005.

[17] R. del Campo, M.-I. Morosini, E. Gómez-G. de la Pedrosa, A. Fenoll, C. Muñoz-Almagro, L. Máiz, F. Baquero, R. Cantón, and the Spanish Pneumococcal Infection Study Network. Population structure, antimicrobial resistance, and mutation frequencies of Streptococcus pneumoniae isolates from cystic fibrosis patients. Journal of Clinical Microbiology, 43:2207–2214, 2005.

[18] E. Denamur, S. Bonacorsi, A. Giraud, P. Duriez, F. Hilali, C. Amorin, E. Bingen, A. Andremont, B. Picard, F. Taddei, and I. Matic. High frequency of mutator strains among human uropathogenic Escherichia coli isolates. Journal of Bacteriology, 184:605–609, 2002.

[19] E. Denamur and I. Matic. Evolution of mutation rates in bacteria. Molecular Microbiology, 60:820–827, 2006.

[20] M. M. Desai and D. S. Fisher. The balance between mutators and nonmutators in asexual populations. Genetics, 188:997–1014, 2011.

[21] J. W. Drake. General antimutators are improbable. Journal of Molecular Biology, 229:8–13, 1993.

[22] J. W. Drake. Too many mutants with multiple mutations. Critical Reviews in Biochemistry and Molecular Biology, 42:247–258, 2007.

[23] J. W. Drake, A. Bebenek, G. E. Kissling, and S. Peddada. Clusters of mutations from transient hypermutability. PNAS, 102:12849–12854, 2005.

[24] J. W. Drake, B. Charlesworth, D. Charlesworth, and J. F. Crow. Rates of spontaneous mutation. Genetics, 148:1667–1686, 1998.

[25] J. W. Drake and J. J. Holland. Mutation rates among RNA viruses. PNAS, 96:13910–13913, 1999.

[26] S. Duffy, L. A. Shackelton, and E. C. Holmes. Rates of evolutionary change in viruses: patterns and determinants. Nature Reviews Genetics, 9:267–276, 2008.

[27] M. Elez, A. W. Murray, L.-J. Bi, X.-E. Zhang, I. Matic, and M. Radman. Seeing mutations in living cells. Current Biology, 20:1432–1437, 2010.

[28] V. Furió, A. Moya, and R. Sanjuán. The cost of replication fidelity in an RNA virus. PNAS, 102:10233–10237, 2005.

[29] M. Furusawa and H. Doi. Promotion of evolution: disparity in the frequency of strand-specific misreading between the lagging and leading DNA strands enhances disproportionate accumulation of mutations. Journal of Theoretical Biology, 157:127–133, 1992.

[30] S. H. Gillespie, S. Basu, A. L. Dickens, D. M. O’Sullivan, and T. D. McHugh. Effect of subinhibitory concentrations of ciprofloxacin on Mycobacterium fortuitum mutation rates. Journal of Antimicrobial Chemotherapy, 56:344–348, 2005.

[31] R. J. Gillies, D. Verduzco, and R. A. Gatenby. Evolutionary dynamics of carcinogenesis and why targeted therapy does not work. Nature Reviews Cancer, 12:487–493, 2012.

[32] D. E. Goldberg, R. F. Siliciano, and W. R. Jacobs Jr. Outwitting evolution: fighting drug-resistant TB, malaria, and HIV. Cell, 148:1271–1283, 2012.

[33] C. Gonzalez, L. Hadany, R. G. Ponder, M. Price, P. J. Hastings, and S. M. Rosenberg. Mutability and importance of a hypermutable cell subpopulation that produces stress-induced mutants in Escherichia coli. PLoS Genetics, 4:e1000208, 2008.

[34] J. Haigh. The accumulation of deleterious genes in a population - Muller’s Ratchet. Theoretical Population Biology, 14:251–267, 1978.

[35] T. Johnson. The approach to mutation-selection balance in an infinite asexual population, and the evolution of mutation rates. Proceedings of the Royal Society B, 266:2389–2397, 1999.

[36] J. M. Kang, N. M. Iovine, and M. J. Blaser. A paradigm for direct stress-induced mutation in prokaryotes. The FASEB Journal, 20:2476–2485, 2006.

[37] S. R. Kennedy, E. M. Schultz, T. M. Chappell, B. Kohrn, G. M. Knowels, and A. J. Herr. Volatility of mutator phenotypes at single cell resolution. PLoS Genetics, 11:e1005151, 2015.

[38] M. Kimura and T. Maruyama. The mutational load with epistatic gene interactions in fitness. Genetics, 54:1337–1351, 1966.

[39] A.G. Knudson. Two genetic hits (more or less) to cancer. Nature Reviews Cancer, 1:157–162, 2001.

[40] M. A. Kohanski, M. A. DePristo, and J. J. Collins. Sublethal antibiotic treatment leads to multidrug resistance via radical-induced mutagenesis. Molecular Cell, 37:311–320, 2010.

[41] N. L. Komarova and D. Wodarz. Drug resistance in cancer: principles of emergence and prevention. PNAS, 102:9714–9719, 2005.

[42] J. E. LeClerc, B. Li, W. L. Payne, and T. A. Cebula. High mutation frequencies among Escherichia coli and Salmonella pathogens. Science, 274:1208–1211, 1996.

[43] H. Lee, E. Popodi, H. Tang, and P. L. Foster. Rate and molecular spectrum of spontaneous mutations in the bacterium Escherichia coli as determined by whole-genome sequencing. PNAS, 109:E2774–E2783, 2012.

[44] C. Lengauer, K. W. Kinzler, and B. Vogelstein. Genetic instabilities in human cancers. Nature, 396:643–649, 1998.

[45] K. R. Loeb and L. A. Loeb. Significance of multiple mutations in cancer. Carcinogenesis, 21:379–385, 2000.

[46] L. A. Loeb. Mutator phenotype may be required for multistage carcinogenesis. Cancer Research, 51:3075–3079, 1991.

[47] L. A. Loeb. A mutator phenotype in cancer. Cancer Research, 61:3230–3239, 2001.

[48] L. A. Loeb, K. R. Loeb, and J. P. Anderson. Multiple mutations and cancer. PNAS, 100:776–781, 2003.

[49] L. A. Loeb, C. F. Springgate, and N. Battula. Errors in DNA replication as a basis of malignant changes. Cancer Research, 34:2311–2321, 1974.

[50] M. Lynch and A. Abegg. The rate of establishment of complex adaptations. Molecular Biology and Evolution, 27:1404–1414, 2010.

[51] R. C. MacLean, C. Torres-Barcelo, and R. Moxon. Evaluating evolutionary models of stress-induced mutagenesis in bacteria. Nature Reviews Genetics, 14:221–227, 2013.

[52] L. M. Mansky and K. S. Cunningham. Virus mutators and antimutators: roles in evolution, pathogenesis and emergence. Trends in Genetics, 16:512–517, 2000.

[53] L. M. Mansky, E. Le Rouzic, S. Benichou, and L. C. Gajary. Influence of reverse transcriptase variants, drugs, and Vpr on Human Immunodeficiency Virus Type 1 mutant frequencies. Journal of Virology, 77:2071–2080, 2003.

[54] L. M. Mansky and H. M. Temin. Lower in vivo mutation rate of human immunodeficiency virus type 1 than that predicted from the fidelity of purified reverse transcriptase. Journal of Virology, 69:5087–5094, 1995.

[55] E. F. Mao, L. Lane, J. Lee, and J. H. Miller. Proliferation of mutators in a cell population. Journal of Bacteriology, 179:417–422, 1997.

[56] I. Matic, M. Radman, F. Taddei, B. Picard, C. Doit, E. Bingen, E. Denamur, and J. Elion. Highly variable mutation rates in commensal and pathogenic Escherichia coli. Science, 277:1833–1834, 1997.

[57] J. D. McCool, E. Long, J. F. Petrosino, H. A. Sandler, S. M. Rosenberg, and S. J. Sandler. Measurement of SOS expression in individual Escherichia coli K-12 cells using fluorescence microscopy. Molecular Microbiology, 53:1343–1357, 2004.

[58] J. H. Miller. Spontaneous mutators in bacteria: insights into pathways of mutagenesis and repair. Annual Reviews in Microbiology, 50:625–643, 1996.

[59] M.-I. Morosini, M.-R. Baquero, J. M. Sanchez-Romero, M.-C. Negri, J.-C. Galan, R. del Campo, J. C. Perez-Diaz, and F. Baquero. Frequency of mutation to rifampicin resistance in Streptococcus pneumoniae clinical strains: hexA and hexB polymorphisms do not account for hypermutation. Antimicrobial Agents and Chemotherapy, 47:1464–1467, 2003.

[60] N. E. Navin. The first five years of single-cell cancer genomics and beyond. Genome Research, 25:1499–1507, 2015.

[61] R. W. Ness, A. D. Morgan, R. B. Vasanthakrishnan, N. Colegrave, and P. D. Keightley. Extensive de novo mutation rate variation between individuals and across the genome of Chlamydomonas reinhardtii. Genome Research, 2015. Published in advance online 10 Aug. 2015.

[62] J. Ninio. Transient mutators: a semiquantitative analysis of the influence of translation and transcription errors on mutation rates. Genetics, 129:957–962, 1991.

[63] A. Oliver, R. Cantón, P. Campo, F. Báquero, and J. Blazquez. High frequency of hypermutable Pseudomonas aeruginosa in cystic fibrosis lung infection. Science, 288:1251–1253, 2000.

[64] A.-L. Prunier, B. Malbruny, M. Laurans, J. Brouard, J.-F. Duhamel, and R. Leclercq. High rate of macrolide resistance in Staphylococcus aureus strains from patients with cystic fibrosis reveals high proportions of hypermutable strains. Journal of Infectious Diseases, 187:1709–1716, 2003.

[65] Y. Ram and L. Hadany. The evolution of stress-induced hypermutation in asexual populations. Evolution, 66:2315–2328, 2012.

[66] Y. Ram and L. Hadany. Stress-induced mutagenesis and complex adaptation. Pro-ceedings of the Royal Society B, 281:2014–1025, 2014.

[67] R. Ribeiro and S. Bonhoeffer. Production of resistant HIV mutants during antiretro-viral therapy. PNAS, 97:7681–7686, 2000.

[68] A. R. Richardson, Z. Yu, T. Popovic, and I. Stojiljkovic. Mutator clones of Neisseria meningitidis in epidemic serogroup A disease. PNAS, 99:6103–6107, 2002.

[69] W. A. Rosche and P. L. Foster. Determining mutation rates in bacterial populations. Methods, 20:4–17, 2000.

[70] S. M. Rosenberg. Spontaneous mutation: real-time in living cells. Current Biology, 20:R810–R811, 2010.

[71] R. Sanjuán, M. R. Nebot, N. Chirico, L. M. Mansky, and R. Belshaw. Viral mutation rates. Journal of Virology, 84:9733–9748, 2010.

[72] P. D. Sniegowski, P. J. Gerrish, T. Johnson, and A. Shaver. The evolution of mutation rates: separating causes from consequences. BioEssays, 22:1057–1066, 2000.

[73] P. D. Sniegowski, P. J. Gerrish, and R. E. Lenski. Evolution of high mutation rates in experimental populations of E. coli. Nature, 387:703–705, 1997.

[74] P. Suárez, J. Valcárcel, and J. Ortin. Heterogeneity of the mutation rates of Influenza A viruses: isolation of mutator mutants. Journal of Virology, 66:2491–2494, 1992.

[75] F. Taddei, M. Radman, J. Maynard-Smith, B. Toupance, P. H. Gouyon, and B. Godelle. Role of mutator alleles in adaptive evolution. Nature, 387:700–702, 1997.

[76] O. Tenaillon, E. Denamur, and I. Matic. Evolutionary significance of stress-induced mutagenesis in bacteria. Trends in Microbiology, 12:264–270, 2004.

[77] C. Torres-Barcelo, M. Kojadinovic, R. Moxon, and R. C. MacLean. The SOS response increases bacterial fitness, but not evolvability, under a sublethal dose of antibiotic. Proceedings of the Royal Society B, 282:20150885, 2015.

[78] M. E. Watson Jr, J. L. Burns, and A. L. Smith. Hypermutable Haemophilus influenzae with mutations in mutS are found in cystic fibrosis sputum. Microbiology, 150:2947–2958, 2004.

[79] C. Zong, S. Lu, A. R. Chapman, and X. S. Xie. Genome-wide detection of single-nucleotide and copy-number variations of a single human cell. Science, 338:1622–1626, 2012.

